# Implicit and explicit learning in reactive and voluntary saccade adaptation

**DOI:** 10.1101/396150

**Authors:** D.M. van Es, T. Knapen

## Abstract

Saccades can either be elicited automatically by salient peripheral stimuli or can additionally depend on explicit cognitive goals. Similarly, it is thought that motor adaptation is driven by the combination of a more automatic, implicit process and a more explicit, cognitive process. However, the degree to which such implicit and explicit learning contribute to the adaptation of more reactive and voluntary saccades remains elusive. To study this question, we employed a global saccadic adaptation paradigm with both increasing and decreasing saccade amplitudes. We assessed the resulting adaptation using a dual state model of motor adaptation. This model decomposes learning into a fast and slow process, which are thought to constitute the explicit and implicit learning, respectively. Our results show that adaptation of reactive saccades is equally driven by fast and slow learning, while fast learning is nearly absent when adapting voluntary (i.e. scanning) saccades. This pattern of results was present both when saccade gain was increased or decreased. Our results suggest that the increased cognitive demands associated with voluntary compared to reactive saccade planning interfere specifically with explicit learning.

## Introduction

Successful interaction with the environment requires that motor actions precisely reach their intended targets. This poses a particularly challenging problem to the organism as conditions both within the body (e.g. waning or gaining of strength) and in the outside world (e.g. wind direction) are in constant flux. The problem is solved by ongoing calibration processes, referred to as motor adaptation. Studying the mechanisms involved in this continuous learning process is achieved by systematically influencing feedback to movements, such as saccades. Saccade adaptation can be induced by displacing the saccade target either closer to (leading to a decrease in saccade gain during adaptation) or further away from (leading to an increase in saccade gain) initial fixation during the executing of the saccade [1].

When saccades are evoked by sudden onset peripheral stimuli, they are relatively fast and commonly referred to as reactive [2]. Alternatively, when saccades depend on behavioral goals they are relatively slow and often termed voluntary [3]. Reactive saccades mainly rely on subcortical structures such as the superior colliculus (SC) and cerebellum as well as on visual and parietal cortical areas (IPS; for comprehensive reviews about the neural structures involved in the generation of the different types of saccades see [4–7]). More complex visual situations may require reactive saccades to be inhibited, while visual information is more thoroughly analyzed in reference to behavioral goals. In this situation, information is relayed to areas in frontal cortex (frontal eye fields, FEF; dorsolateral prefrontal cortex, DLPFC; anterior cingulate cortex, ACC; [8]). FEF and IPS can then project directly back to the SC [9,10], to generate such voluntary saccades.

The adaptation of voluntary and reactive saccades is similarly thought to depend on differential cortical and subcortical contributions [3]. The first evidence for separate mechanisms involved in reactive and voluntary saccades comes from findings of asymmetrical transfer of adaptation between saccade types [2,11,12]. Additionally, it was shown that voluntary and reactive saccades can be adapted simultaneously in different directions [13] and that adaptation of voluntary but not reactive saccades transferred to hand pointing motions [14]. These and other findings [11] have led to the suggestion that the cerebellum and subcortical structures are important for adaptation of both types of saccades, while adaptation of voluntary saccades additionally depends on upstream areas [3]. These upstream areas likely constitute the fronto-parietal areas involved in the generation of voluntary saccades [15,16]. In sum, both the generation and adaptation of voluntary saccades place greater demands on fronto-parietal areas.

These differential neural contributions in voluntary and reactive saccades can be well understood in as differential cognitive involvement in terms of attention and working memory [17]. Although the exact functional link between attention and saccades remains elusive, different models of attention suggest a close relation [18–20]. Voluntary saccades especially demand the selective enhancement and suppression of visual information. In addition, voluntary saccades require the active maintenance of internalized stimulus-response rules in working memory. Indeed, voluntary saccade performance degrades with increasing working memory load [21,22], and depends on individual differences in working memory capacity [23,24]. Moreover, the network of fronto-parietal areas that contributes to voluntary saccades (IPS, FEF, DLPFC) closely overlaps with those associated with attention [25–29] and working memory [30,31].

Saccadic adaptation already occurs within the first few perturbation trials and continues throughout an experimental block. This learning also carries over to subsequent learning blocks, producing savings (i.e. quicker relearning), interference (i.e. slower learning of opposing adaptation) and spontaneous recovery (i.e. rebound of previous learning after forgetting [32,33]). These and other phenomena of motor learning can be well accounted for by a dual-state model of adaptation [34,35]. This model posits that learning is driven (1) by a fast process that learns and forgets quickly and (2) by a slow process that learns and forgets slowly. Recent studies have suggested that the fast process reflects the explicit and voluntary effort to actively counteract the perturbation (i.e. ‘one-shot’ learning), whereas the slow process reflects implicit and automatic learning [36,37].

While a cerebellum-dependent automatic learning process has been well established [38–40], it has been suggested that the explicit component of learning should be viewed in terms of cognitive processes [41,42]. In fact, it was shown that working memory capacity correlates with explicit and not implicit learning performance (in motor sequence learning [43], and in visuomotor adaptation [44]). In addition, it was shown that an interfering attentional task reduced the overall amount of learning during visuomotor adaptation [45], especially early during learning [46]. Moreover, early visuomotor adaptation particularly induced fronto-parietal activations [47–49], which were in turn shown to be related to spatial working memory [44,49].

Taken together, the studies described above reveal striking similarities between the mechanisms involved in explicit learning and voluntary saccades and between those involved in implicit learning and reactive saccades. Specifically, both implicit learning and reactive saccades depend predominantly on automatic processes mediated by cerebellum, whereas explicit learning and voluntary saccades additionally require spatial cognition mediated by fronto-parietal areas. This raises the question to what extent implicit and explicit learning contribute to reactive and voluntary saccades. On the one hand, it is possible that the type of learning is contingent upon the neural structures involved in generation of the saccade. This would mean that reactive saccade adaptation elicits relatively stronger implicit learning (i.e. both cerebellum dependent), whereas voluntary saccade adaptation elicits relatively stronger explicit learning (as both depend on fronto-parietal regions). On the other hand, it is possible that the increased cognitive load associated with voluntary saccades curtails the remaining capacity available for explicit learning. This would be in line with cognitive load theories of learning [50,51].

Furthermore, different mechanisms have been posited for gain-up and gain-down adaptation [32,52–56]. These differences are thought to be in cerebellar but not cortical contributions [57–61]. Therefore, we expect no differential contribution of implicit and explicit learning to gain-down and gain-up adaptation.

To test these hypotheses, we employed a global saccadic adaptation paradigm [62]. During the execution of the saccade, the target was displaced either further away from (gain-up adaptation) or closer to (gain-down adaptation) previous fixation. Saccades were either elicited exogenously by a peripheral cue (reactive saccades), or endogenously by means of an internalized instruction to move to one of multiple targets (scanning saccades, a type of voluntary saccade). The path traveled across saccades was kept constant between the different saccade conditions. In addition, we hypothesized that awareness of the target displacement should increase the amount of explicit learning. To investigate this, we asked participants on each trial whether they perceived the target displacement. We assessed contributions of implicit and explicit learning as the gain of the slow and fast state respectively, as derived from the dual-state model of adaptation [34].

## Methods

### Participants

Twelve participants (5 female) participated in this study. All participants gave written informed consent for participation. The study was approved by the ethical committee of the Vrije Universiteit Amsterdam.

### Apparatus

Stimuli were presented on a CRT monitor with a resolution of 1024×768 at a vertical refresh rate of 120Hz. Eye movements were recorded at 1000Hz using an Eyelink 1000 Tower Mount (SR Research, Osgoode, Ontario, Canada). A 9-point calibration procedure was run before the start of each experimental condition. All light sources were eliminated in the experimentation room in order to minimize visual references. To eliminate remaining visible screen edges, we placed a red filter over the screen, only allowing red to pass through (+/- > 650 nm; Lee Filters color #787 ‘Marius Red’). All stimuli were consequently presented using only the red CRT channel. Thus, participants were in complete darkness and only viewed red stimuli.

### Task

Participants followed a square target (0.5×0.5 dva) around the screen along a clockwise hexagonal path, as depicted in Fig 1A. Participants started each block fixating at the central and lower position. On all trials in all conditions, the currently fixated target flashed for 150 ms to indicate the eye movement instruction. In the reactive saccade condition the current fixation target extinguished and immediately appeared in the next location along the hexagonal, provoking a peripheral visual transient. In the scanning saccade condition all possible locations along the hexagonal were always present on the screen, meaning there were no peripheral visual transients before the saccade. Importantly, in both saccade conditions participants completed identical saccade paths. The only difference was the number of presented targets (i.e. only one or all). This ensured that in the reactive saccade conditions subjects could simply follow the target around the screen, whereas in the scanning saccade conditions participants had to actively select one of multiple targets and plan a saccade to it. As soon as the initial saccade was detected by experimental software, the target(s) displaced either 3.3 dva further along the saccade trajectory (gain-up adaptation), or 2.5 dva back in the opposite direction of the saccade trajectory (gain-down adaptation). These differential absolute displacement magnitudes ensured equal relative gains (7.5/10 ≈ 10/13.3). Thus, in the reactive saccade condition only the single target was displaced, whereas in the scanning condition all targets displaced simultaneously.

**Fig 1.**
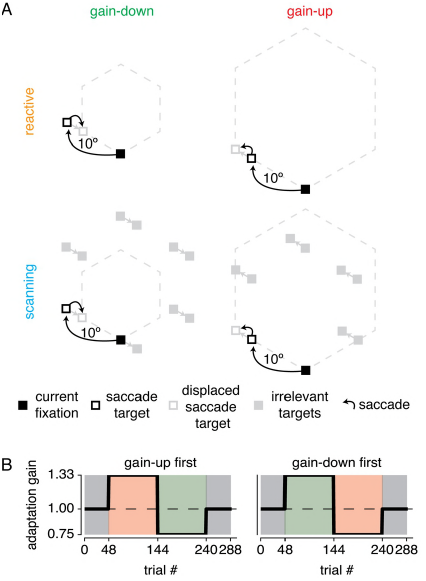
Experimental design. (A) Participants made saccades to targets around a clockwise hexagonal path. In the reactive saccade condition, a single target sequentially jumped along the corners of the hexagonal. Subjects were instructed to simply follow the target. In the scanning saccade condition, all targets were always presented and subjects were instructed to saccade to the next clockwise target when the currently fixated target flashed. During the saccade, targets jumped either further away from (gain-up) or closer to (gain-down) previous fixation. Participants sat in full darkness and all stimuli were filled red squares. Note that dotted outline and different appearance of squares are for illustrative purposes only. (B) The experiment was made up of an initial baseline block, two adaptation blocks and a final baseline block. All participants completed two experimental sessions per saccade type, switching the adaptation block orders between gain-up and gain-down.

To ensure unpredictability of the saccade target onset time, the fixation interval was randomly selected from an exponential distribution with minimum of 250 ms and mean of 550 ms. Subsequently, the currently fixated target flashed (150 ms) after which polling for saccades started (using threshold of 1.5 dva; maximally 2750 ms). As soon as the saccade was detected, the saccade target was displaced.

To asses awareness of the target displacement, participants indicated on each trial whether they perceived the target displacement. For this, the square saccade target turned into a triangle that pointed either left- or rightwards, 500 ms after saccade target displacement. Participants were instructed to press in the direction of the triangle to report the displacement as ‘seen’ and in the opposite direction to indicate ‘unseen’. The response window ended as soon as a response was given or when 2000 ms passed.No feedback was provided.

### Procedure

Each experimental condition was made up of four blocks (see Fig 1B). In the first and last baseline blocks of 48 trials no target displacements took place to establish baseline measurements of saccade parameters. The second and third blocks contained 96 trials of either gain-up or gain-down adaptation. During the course of an adaptation block, participants learned the regularity in the target displacement and gradually adapted the gain of their initial saccade to land closer to the post-displacement location. In total, there were four experimental sessions (i.e. full factorial design with factors saccade type [reactive/scanning] and adaptation direction order [up-down/down-up]). The order of the four conditions were randomized on the factor of saccade type, yielding 12 possible orders (4!/2) each assigned to a different participant. Each experimental session lasted approximately 10 minutes. To ensure that adaptation had fully returned to baseline, the different experimental sessions were separated by at least 2 hours. Before the first experimental session, participants practiced a shortened version of the experiment (6 saccades per block). This allowed them to estimate expected jump magnitude to judge as ‘seen’ or ‘unseen’ and to practice with the task procedure in general.

### Data analysis

#### Saccade amplitude definition

Eyetracking data were preprocessed using the hedfpy package (https://github.com/tknapen/hedfpy). Offline saccade detection was performed using the Engbert and Mergenthaler algorithm [63]. The preprocessed data and the analyses presented in this manuscript can be found under https://figshare.com/s/2d97ad68b6ec3801314c and https://figshare.com/s/eba20e1e506b4f4d8cf2 respectively. Saccade amplitudes were converted to ‘gain’ by dividing them by the median saccade amplitude from the first baseline block, for each of the 6 hexagonal directions separately. Trials were rejected when (1) it included a blink, (2) saccade amplitude was below 3 dva or more than 3 two-sided median absolute deviations in that block and (3) saccade start point deviated more than 1.5 dva from the pre-saccadic target.

#### Exponential fits to first block

In order to quantify the timescale of adaptation in the first block we fitted exponentials of the form:

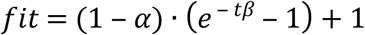

where *α* and *β* reflect a gain and timescale parameter respectively and *t* reflects trial number. This form of the exponential equation ensures that it starts at 1 and approaches (1 − *α*) at infinite *t*. Including the gain parameter (*α*) ensures that the estimate of the timescale (*β*) is unaffected by overall adaptation magnitude. Estimates for the reliability of parameters in this function were found by using a 10^5^ fold bootstrap procedure. Within each fold of this procedure, the exponential function was fitted using least-squares to average data of a particular condition across a random sample of participants with replacement (cf. Fig A,B). In order to retrieve differences between conditions, exponential functions were fitted to all relevant conditions within each fold, ensuring paired comparisons between bootstrap samples of participants (cf. Fig C). Resulting p-values were calculated as the ratio of parameter difference estimates that fell below versus above 0, multiplied by 2 (i.e. two-tailed tests).

#### Fast and slow process contribution

In order to determine the fast and slow process contributions to reactive and scanning saccade adaptation, we fitted the multi-rate model by [34] to the data. We set learning and retention parameters to the values established in the original study (fast learn = .21, fast retention = .59, slow learn = .02 and slow retention = .992), and varied gain parameters that scaled the contribution of each process. We fixed the learning and retention parameters in order to maximize stability of our parameters of interest (i.e. the gain parameters). Thus, adaptation was given by the following:

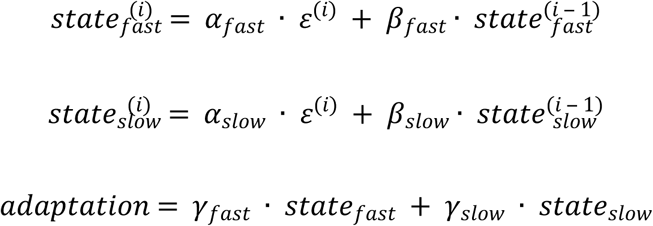

where *α*, *β*, γ, *∈* and *i* refer to learning rate, retention rate, process gain, saccade error and trial index respectively.

We also fitted the model using freely varying learning and forgetting parameters. This yielded similar results, although parameter estimates were less stable. This is likely caused by the fact that parameter estimates trade off in the fitting procedure. This interaction between model parameters is illustrated by the following. When learning is relatively strong and forgetting relatively weak, this results in greater overall learning. Similarly, increasing the gain parameter while keeping the learning and forgetting rates equal also results in greater overall learning. Although this latter approach fixes the shape of the individual slow and fast processes, it does allow for the shape of overall learning to vary as a result of differential weighing of the fast and slow processes. As we are specifically interested in the shape of the overall learning curve and not in the shape of either the fast or slow processes alone, we opted to fix the learning and forgetting parameters in order to maximize the stability of the gain parameters. Future studies interested in differences in the shape of fast and slow learning could include error-clamp trials, where visual error is eliminated by displacing the saccade target to the saccade endpoint [35].

## Results

First, we verified whether we successfully evoked reactive and voluntary saccades by investigating their relative latencies. Reactive saccade latency is usually around 200 ms, whereas voluntary saccades latency commonly exceeds 250 ms [3]. Our results indeed showed that reactive compared to scanning saccades were faster (193 vs. 307 ms; see Fig 2; F_(1,11)_ = 34.348, p = 1.092*10 ^−4^, η^2^p= .833). In addition, saccade latency was not different for gain-up vs. gain down adaptation (F_(1,11)_ = 1.010, p = .337, η^2^p= .007), nor was there an interaction between saccade type and adaptation direction (F_(1,11)_ = 0.292, p = .600, η^2^p= .003)).

**Fig 2.**
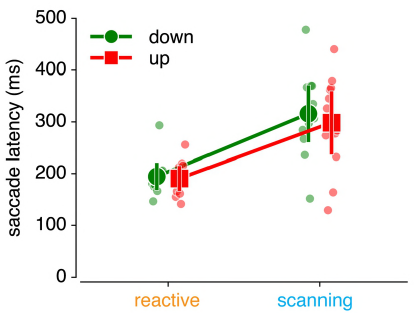
Saccade latencies. Saccade latencies in the second block of each saccade adaptation condition show that scanning saccades are slower than reactive saccades.

Fig 3 shows saccade amplitude across trials and highlights several features. First, and in correspondence with the literature, it shows stronger gain-down compared to gain-up adaptation [64]. In addition, it shows that adaptation of scanning and reactive saccades is of comparable magnitude. Second, it suggests that reactive saccade adaptation reaches maximal adaptation early in the block, whereas scanning saccade adaptation continues more strongly throughout the block. Finally, adaptation seems to change more strongly between blocks 2 and 3 in the reactive compared to the scanning saccade conditions.

**Fig 3.**
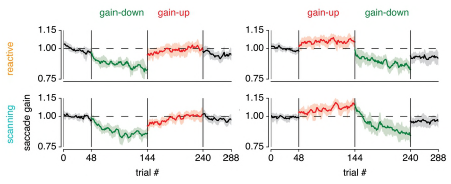
Saccade adaptation time-courses for the different conditions. Saccade amplitude across trials for the different saccade conditions (reactive and scanning) and for the different orders of gain direction blocks (down-up or up-down). Saccade amplitude is depicted as the ratio to median saccade amplitude in the first block (per saccade direction, see Methods) and shown as running average over 6 saccades (the hexagonal period). Shaded areas correspond to 95% CI over participants.

To quantify these observations, we first analyzed the timescale of adaptation. Fig 4 shows an exponential function fitted to the data from the second block (i.e. the first adaptation block), both for gain-up and gain-down adaptation. Indeed, the timescale of this exponential was slower for scanning compared to reactive saccades, both in the gain-down (p = .007) and gain-up (p = .006) blocks (Fig 4A and B). Furthermore, we tested whether adaptation changed more between the second and the third block. The magnitude of such a sudden change indicates the combination of two processes: (1) the speed of forgetting of adaptation of the previous block and (2) the speed of learning at the beginning of the new block. Confirming the visual intuition described above, we find greater changes in adaptation state between the second and third block in the reactive compared to the scanning condition, for both gain-direction reversals (quantified as the mean gain over the last and first 6 trials of each block; down-up t_(11)_ = 2.892, p = 0.015, Cohen‘s d = 0.872, up-down t_(10)_ = -3.099, p = 0.011, Cohen’s d = -0.980, see Fig 4D; note that one subject was not included in this latter analysis as he/she did not happen to have any valid saccades in the trials analyzed; see Methods for saccade exclusion criteria). This implies that the speed of forgetting and/or the speed of learning is faster in reactive compared to scanning saccades. Together, these results show that reactive compared to scanning saccade adaptation occurs at a faster timescale.

**Fig 4.**
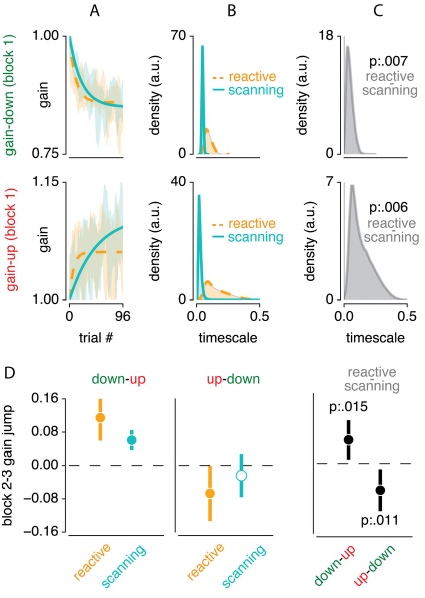
Indications for fast and slow process contributions. (A) Exponential functions fitted to the first adaptation block of each condition. (B) Bootstrapped distribution of the timescale parameter across participants, for the different conditions. (C) Difference in this timescale between reactive and scanning for both gain direction conditions. This shows that reactive compared to scanning saccade adaptation occurs at a faster rate. (D) Changes in saccade gain between the average of the last 6 saccades of the second block and the average of the first 6 saccades of the third block. Results are shown for both gain direction reversals (up-to-down and down-to-up) for both reactive and scanning saccades, and for the difference between these saccade types. This shows that when adapting reactive compared to scanning saccades, adaptation changes more quickly from one to another gain-change situation.

We next investigated whether the faster timescale observed above can be explained in terms of dual state decomposition into a fast and slow process [34]. For these analyses, we focused on data from the second block (i.e. first adaptation block). Fig 5 shows the fitted slow and fast processes to the data. Visual inspection of this figure suggests that the slow process contributes more to overall adaptation in scanning compared to reactive saccades.

**Fig 5.**
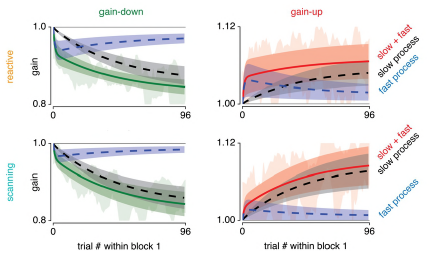
Dual-rate model fits to gain-up and gain-down adapted reactive and scanning saccades. Fitted fast (blue) and slow (black) processes to overall adaptation (green when gain-down and red when gain-up) to data from the second block. Shaded areas around model fits represent the 95% CI over participants. Underlying shaded data area indicates 95% CI over participants of moving average over six trials (as in Fig 3).

The gains of both processes are summarized in Fig 6A. Overall process gain was roughly equal between reactive and scanning saccades (0.576 vs 0.509 respectively, main effect of condition F_(1,11)_ = 3.810, p = .077, η^2^p = .013). Adaptation gain was indeed much higher for gain down compared to gain up adaptation (0.784 vs 0.301 respectively, main effect of direction F_(1,11)_ = 40.342, p = 5.434 * 10 ^−5^, η^2^p = .650). Yet, this difference was not different between reactive and scanning saccades (interaction between condition and direction, F_(1,11)_ = 1.771, p = .210, η^2^p= .008). Also, adaptation was driven more strongly by the slow compared to the fast process (0.665 vs 0.420 respectively, main effect of process F_(1,11)_ = 12.869, p = 0.004, η^2^p = .167). This difference was mainly driven by gain-down as opposed to gain-up adaptation (0.396 vs 0.094 respectively, interaction beween direction and process F_(1,11)_ = 6.076, p = .031, η^2^ p = .063). However, this interaction could be driven by increased overall adaptation magnitude in gain-down compared to gain-up adaptation. To account for this, we calculated the ratio of slow compared to overall (fast+slow) gain and compared this ratio between gain-up and gain-down adaptation. This showed that when controlling for overall adaptation gain, there is no differential contribution of the fast and slow processes between gain-up and gain-down adaptation (p = .667). Of particular importance to the purpose of this study, the fast and slow process contributed differently to reactive and scanning saccades (interaction between condition and process, F_(1,11)_ = 10.256, p = .008, η^2^p = .114). This differential contribution of the fast and slow process to reactive and scanning saccades was not different between gain-up/gain-down adaptation (three-way interaction between condition, direction and process, F_(1,11)_ = 0.160, p = .696, η^2^p = .002).

**Fig 6.**
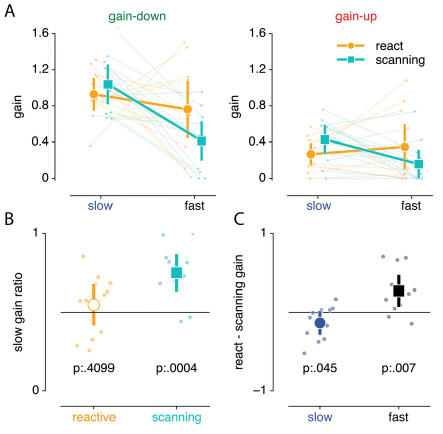
Quantification of dual process contribution to reactive and scanning saccades. (A) Comparison of process gain in both saccade type (reactive and scanning) and direction (gain-up and gain-down) blocks. (B) Contribution of slow process to overall adaptation averaged over gain-down and gain-up blocks for reactive and scanning blocks separately. This shows that reactive saccades are equally driven by the fast and slow process, whereas scanning saccades are driven mainly by the slow process. (C) Reactive minus scanning process gain averaged over gain-up and gain-down blocks for the slow and fast process separately. This shows that the slow process is stronger in scanning compared to reactive saccades, while the fast process is stronger in reactive compared to scanning saccades. Error bars depict 95% CI over participants. Individual dots and lines indicate individual participants.

The main purpose of this study was to establish whether the fast or slow process contributes more to both scanning and reactive saccade adaptation. In order to investigate this, we further explored the differential contribution of the fast and slow process to overall adaptation. Since this was not different between gain-up and gain-down adaptation, we averaged over both gain direction conditions. We first analyzed the ratio of slow compared to overall (slow+fast) gain (Fig 6B). This showed that the slow and fast process contributed equally to reactive saccade adaptation (mean slow ratio of 0.547 was not different from 0.5 with t_(11)_ = 0.857, p = .410, Cohen‘s d = 0.258), while scanning saccade adaptation was mainly driven by the slow process (mean slow ratio of 0.749 was different from 0.5 with t_(11)_ = 5.053, p = 3.705 * 10 ^−4^, Cohen’s d = 1.523). This differential process contribution could either be due to increased slow or decreased fast process contribution in the scanning compared to reactive saccade adaptation. To investigate this, we computed the difference in adaptation gain between the reactive and scanning conditions for both fast and the slow process (Fig 6C). This showed that slow process gain is 0.135 larger in scanning compared to reactive saccade adaptation (t_(11)_ = 2.265, p = .045, Cohen’s d = 0.683). Conversely, fast process gain was 0.270 larger in the reactive compared to the scanning saccade conditions (t_(11)_ = 3.267, p = .007, Cohen’s d = 0.985).

In sum, these results show that scanning saccade adaptation was driven relatively more by the slow compared to the fast process, whereas reactive saccade adaptation is driven equally by fast and slow process. The difference between reactive and scanning saccade adaptation is mainly due to a decreased fast process in scanning saccades, but also by a slight increase in the slow process in reactive saccades.

In addition to measuring saccade accuracy, we asked subjects to report their awareness of the target displacement in a binary fashion (i.e. seen or not seen; see Methods). Fig 7A depicts this seen judgement throughout the experiment. First, this confirms that subjects generally did not report any displacements in the baseline blocks. Second, it shows that subjects saw the displacement most strongly at the beginning of each adaptation block, to then gradually become more invisible. To relate these findings to the results presented above, we further analyzed data from the second block (i.e. first adaptation block). Fig 7B summarizes the average judgements across conditions. This shows that the displacement was seen more in scanning compared to reactive saccade adaptation conditions (F_(1,11)_ = 11.954, p = .005, η^2^p = .124), and more in the gain-down compared to gain-up conditions (F_(1,11)_ = 10.204, p = 0.009, η^2^p= .234). There was no interaction between saccade type and gain direction (F_(1,11)_ = 0.027, p = .872, η^2^p = 2.401* 10 ^−4^).

**Fig 7.**
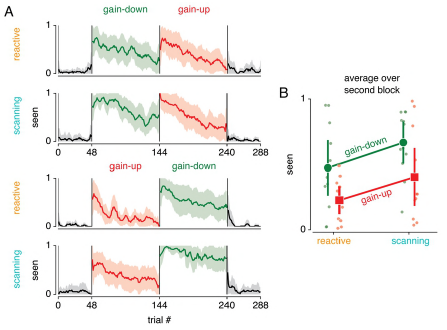
Saccade target displacement awareness. (A) Across participant average of binary displacement awareness (i.e. seen versus unseen) across the experiment. This verifies that the target displacement was not seen in the baseline blocks (i.e. blocks 1 and 4 colored black). In addition, it shows that target displacement was seen more often at the beginning compared to the end of an adaptation block. (B) Average displacements awareness per condition in the second block. The target displacement was perceived more often in gain-down compared to gain-up adaptation, and more often in scanning compared to reactive saccade conditions (see main text for statistics).

## Discussion

We studied the contributions of the fast and slow process [34] to reactive and scanning gain-down and gain-up saccade adaptation. First, we showed that gain-up adaptation was weaker overall compared to gain-down adaptation. Second, our results showed that scanning saccade adaptation was driven more by the slow compared to the fast process, whereas reactive saccade adaptation was driven equally strongly by the fast and slow process. This was caused mainly by an increased fast process contribution but also by a decreased slow process contribution in reactive compared to scanning saccade adaptation. Third, we found no differential fast and slow process contribution to gain-down and gain-up adaptation. Finally, we found that the target displacement was seen more often in scanning compared to reactive saccade adaptation and more often in gain-down compared to gain-up adaptation.

The fast and slow process were recently associated with explicit and implicit learning respectively [36,37]. Implicit learning refers to automatic learning that cannot be actively engaged or disengaged, which is thought to reflect ongoing recalibration of an internal forward model of the motor apparatus. In contrast, explicit learning refers to the voluntary and active effort to flexibly alter motor plans, expressing most strongly early in learning. While implicit learning is thought to be mainly cerebellum dependent, explicit learning likely also requires contributions from frontal cortex [42,44,47–49,65]. In correspondence, reactive saccades are thought to be generated mainly subcortically, whereas voluntary saccades also require the involvement of fronto-parietal areas [3,15,16,66]. We therefore suggested that learning type (i.e. explicit versus implicit) could be contingent upon the neural structures involved in the generation of the movement. This would have predicted that reactive saccade adaptation should mainly be driven by implicit learning while scanning saccade adaptation should mainly be driven by explicit learning. Our results however disconfirm this notion by showing that reactive saccade adaptation was driven equally by slow and fast learning, while scanning saccade adaptation was driven predominantly by slow learning.

The network of areas involved in both explicit learning and voluntary saccades agree well with the multiple demand system, a proposed neural correlate of the cognitive executive system [67]. Voluntary saccades involve processes that can be considered “cognitive”, such as the suppression and enhancement of visual information (i.e. attention) and a comparison to internal behavioral goals (i.e. working memory). In fact, these increased cognitive demands make voluntary saccade performance a suitable tool for characterize cognitive deficits in psychopathology [68]. Yet, increasing cognitive load is known to interfere with learning [50,51]. Specifically, it was shown that explicit and not implicit learning is related to individual differences in working memory capacity [43,44]. In addition, performing simultaneous secondary tasks interfered with motor adaptation by taxing the executive system [45], especially early in learning [46]. The increased cognitive demands associated with voluntary saccades can therefore explain the impaired explicit learning we observed in scanning compared to reactive saccades. In addition, it was previously suggested that increased explicit learning inhibits implicit learning [69]. Conversely, reducing explicit learning should therefore allow for improved implicit learning. Our data seem to support such a trade-off mechanism as we found that both explicit learning was increased and implicit learning was decreased in reactive compared to scanning saccade adaptation. Taken together, this suggests that increasing cognitive load associated with movement execution reduces explicit learning, which in turn leads to improved implicit learning.

This cognitive load [50,51] interpretation of our results is further supported by studies of healthy aging. Such studies generally find that aging is associated with degrading cognitive performance [70], and decreases in frontal lobe volume [71–73]. In addition, aging is related to reductions in explicit but not implicit learning [74–80]. Moreover, these age-related declines in explicit learning were related to the age-related declines in cognitive performance [74,81,82]. Finally, voluntary saccade performance also decreases with increasing age [83–85]. Together, this supports the notion that both voluntary saccades and explicit learning depend on availability of cognitive resources mediated by frontal cortex.

As we expected no differential cognitive load, nor a differential recruitment of frontal cortex between gain-up and gain-down adaptation, we expected no differential implicit and explicit learning contribution in gain-down and gain-up adaptation. Indeed, our data showed that gain-up and gain-down adaptation were not differently driven by explicit and implicit learning. Yet, our data do confirm earlier reports showing that gain-up adaptation results in weaker overall adaptation compared to gain-down adaptation [53,54].

Finally, we hypothesized that increased target jump awareness should lead to increased explicit learning. However, even though target jump awareness was greater in gain- down compared to gain-up adaptation, we found no differential fast learning between these conditions. In addition, even though target jump awareness was greater in scanning compared to reactive saccade adaptation, we found decreased fast learning in scanning compared to reactive saccade adaptation. This suggests that explicit learning did not depend on target jump awareness. Although this is contradictory to our initial hypothesis, it can be explained by a disconnect between definition of the target jump on the one hand, and of the saccade endpoint error on the other hand. Specifically, target jump magnitude is given by the distance between the pre-saccadic and the post- saccadic target. Yet, saccade error magnitude is given by the distance of the post- saccadic target to the saccade landing position. In other words, when a saccade lands exactly at a displaced saccadic target position (i.e. perfect adaptation), there is no saccade error but still a considerable target jump. Conversely, when a saccade lands some distance away from a non-displaced saccadic target, there is considerable saccadic error but no target jump. This disconnect between target jump and saccade error magnitude is further exacerbated by the overrepresentation of the fovea in retinotopically organized areas. While the target jump is differently warped by this foveal magnification depending on the direction of the jump in relation to the direction of the saccade, the saccadic error is always relative to fixation. This means that the retinotopic representation of a target displacement is greater when it jumps towards versus away from the fovea. This can explain why gain-down compared to gain-up adaptation resulted in greater target jump awareness. In fact, the target jumped across the fovea in gain-down adaptation, while it moved further into the periphery in gain-up adaptation. Finally, the greater target jump awareness in scanning compared to reactive saccade adaptation is likely explained by the increased visual references in the scanning saccade condition induced by all targets being presented simultaneously. In sum, this can explain why target jump awareness was unrelated to explicit learning. We suggest that future studies interested in directly measuring the explicit component of saccadic adaptation should therefore gauge explicit knowledge of the saccadic endpoint error rather than awareness of the target jump.

In sum, our results suggest that increasing cognitive load during movement execution reduces the capacity for explicit learning. In general, this suggests that increasingly simplified training environments will enhance performance particularly when little time for learning is available. Conversely, in situations that place great demands on cognitive resources, learning will be mainly based on implicit learning and thus will take time to develop.

## Acknowledgement

We would like to thank Tinka Beemsterboer for her contributions to this project.

## References

1. McLaughlin SC. Parametric adjustment in saccadic eye movements. Percept Psychophys. 1967;2: 359–362.

2. Deubel H. Separate adaptive mechanisms for the control of reactive and volitional saccadic eye movements. Vision Res. 1995;35: 3529–3540.

3. Pelisson D, Alahyane N, Panouillères M, Tilikete C. Sensorimotor adaptation of saccadic eye movements. Neurosci Biobehav Rev. 2010;34: 1103–1120.

4. Pierrot-Deseilligny C, Milea D, Müri RM. Eye movement control by the cerebral cortex. Current Opinion in Neurology. 2004;17: 17.

5. McDowell JE, Dyckman KA, Austin BP, Clementz BA. Neurophysiology and neuroanatomy of reflexive and volitional saccades: Evidence from studies of humans. Brain Cognition. 2008;68: 255–270.

6. Munoz DP, Everling S. Look away: the anti-saccade task and the voluntary control of eye movement. Nat Rev Neurosci. 2004;5: 218–228.

7. Müri RM, Nyffeler T. Neurophysiology and neuroanatomy of reflexive and volitional saccades as revealed by lesion studies with neurological patients and transcranial magnetic stimulation (TMS). Brain Cognition. 2008;68: 284–292.

8. Schall JD. Visuomotor Functions in the Frontal Lobe. Annu Rev Vis Sci. 2015;1: 469–498.

9. Helminski JO, Segraves MA. Macaque Frontal Eye Field Input to Saccade-Related Neurons in the Superior Colliculus. J Neurophysiol. 2003;90: 1046–1062.

10. Baizer JS, Desimone R, Ungerleider LG. Comparison of subcortical connections of inferior temporal and posterior parietal cortex in monkeys. Visual Neurosci. 1993;10: 59–72.

11. Alahyane N, Salemme R, Urquizar C, Cotti J, Guillaume A, Vercher J-L, et al. Oculomotor plasticity: Are mechanisms of adaptation for reactive and voluntary saccades separate? Brain Res. 2007;1135: 107–121.

12. Cotti J, Panouillères M, Munoz DP, Vercher J-L, Pélisson D, Guillaume A. Adaptation of reactive and voluntary saccades: Different patterns of adaptation revealed in the antisaccade task. J Physiol-London. 2009;587: 127–138.

13. Fujita M, Amagai A, Minakawa F, Aoki M. Selective and delay adaptation of human saccades. Brain Res Cogn Brain Res. 2002;13: 41–52.

14. Cotti J, Guillaume A, Alahyane N, Pélisson D, Vercher J-L. Adaptation of voluntary saccades, but not of reactive saccades, transfers to hand pointing movements. J Neurophysiol. 2007;98: 602–612.

15. Panouillères M, Habchi O, Gerardin P, Salemme R, Urquizar C, Farne A, et al. A Role for the parietal cortex in sensorimotor adaptation of saccades. Cereb Cortex. 2014;24: 304–314.

16. Gerardin P, Miquée A, Urquizar C, Pélisson D. Functional activation of the cerebral cortex related to sensorimotor adaptation of reactive and voluntary saccades. NeuroImage. 2012;61: 1100–1112.

17. Hutton SB. Cognitive control of saccadic eye movements. Brain Cognition. 2008;68: 327–340.

18. Rizzolatti G, Riggio L, Dascola I, Umiltá C. Reorienting attention across the horizontal and vertical meridians: Evidence in favor of a premotor theory of attention. Neuropsychologia. 1987;25: 31–40.

19. Schneider WX. VAM: A neuro-cognitive model for visual attention control of segmentation, object recognition, and space-based motor action. Visual Cognition. 1995;2: 331–376.

20. Wollenberg L, Deubel H, Szinte M. Visual attention is not deployed at the endpoint of averaging saccades. PLoS Biol. 2018;16: e2006548.

21. Roberts RJ, Hager LD, Heron C. Prefrontal cognitive processes: Working memory and inhibition in the antisaccade task. J Exp Psychol Gen. 1994;123: 374–393.

22. Mitchell JP, Macrae CN, Gilchrist ID. Working memory and the suppression of reflexive saccades. J Cognitive Neurosci. 2002;14: 95–103.

23. Kane MJ, Bleckley MK, Conway AR, Engle RW. A controlled-attention view of working-memory capacity. J Exp Psychol Gen. 2001;130: 169–183.

24. Unsworth N, Schrock JC, Engle RW. Working memory capacity and the antisaccade task: Individual differences in voluntary saccade control. J Exp Psychol Learn Mem Cogn. 2004;30: 1302–1321.

25. Nobre AC, Gitelman DR, Dias EC, Mesulam MM. Covert visual spatial orienting and saccades: Overlapping neural systems. NeuroImage. 2000;11: 210–216.

26. Beauchamp MS, Petit L, Ellmore TM, Ingeholm J, Haxby JV. A parametric fMRI study of overt and covert shifts of visuospatial attention. NeuroImage. 2001;14: 310–321.

27. Kastner S, DeSimone K, Konen CS, Szczepanski SM, Weiner KS, Schneider KA. Topographic maps in human frontal cortex revealed in memory-guided saccade and spatial working-memory tasks. J Neurophysiol. 2007;97: 3494–3507.

28. Szczepanski SM, Pinsk MA, Douglas MM, Kastner S, Saalmann YB. Functional and structural architecture of the human dorsal frontoparietal attention network. Proc Natl Acad Sci USA. 2013;110: 15806–15811.

29. Corbetta M, Shulman GL. Control of goal-directed and stimulus-driven attention in the brain. Nat Rev Neurosci. 2002;3: 201–215.

30. Sprague TC, Ester EF, Serences JT. Reconstructions of information in visual spatial working memory degrade with memory load. Curr Biol. 2014;24: 2174–2180.

31. Christophel TB, Klink PC, Spitzer B, Roelfsema PR, Haynes J-D. The distributed nature of working memory. Trends Cogn Sci. 2017;21: 111–124.

32. Kojima Y, Iwamoto Y, Yoshida K. Memory of learning facilitates saccadic adaptation in the monkey. J Neurosci. 2004;24: 7531–7539.

33. Miall RC, Jenkinson N, Kulkarni K. Adaptation to rotated visual feedback: A reexamination of motor interference. Exp Brain Res. 2004;154: 201–210.

34. Smith MA, Ghazizadeh A, Shadmehr R. Interacting adaptive processes with different timescales underlie short-term motor learning. PLoS Biol. 2006;4: e179.

35. Ethier V, Zee DS, Shadmehr R. Spontaneous recovery of motor memory during saccade adaptation. J Neurophysiol. 2008;99: 2577–2583.

36. McDougle SD, Bond KM, Taylor JA. Explicit and implicit processes constitute the fast and slow processes of sensorimotor learning. J Neurosci. 2015;35: 9568–9579.

37. Huberdeau DM, Krakauer JW, Haith AM. Dual-process decomposition in human sensorimotor adaptation. Curr Opin Neurobiol. 2015;33: 71–77.

38. Straube A, Deubel H, Ditterich J, Eggert T. Cerebellar lesions impair rapid saccade amplitude adaptation. Neurology. 2001;57: 2105–2108.

39. Synofzik M, Lindner A, Thier P. The cerebellum updates predictions about the visual consequences of one’s behavior. Curr Biol. 2008;18: 814–818.

40. Izawa J, Criscimagna-Hemminger SE, Shadmehr R. Cerebellar contributions to reach adaptation and learning sensory consequences of action. J Neurosci. 2012;32: 4230–4239.

41. Taylor JA, Ivry RB. Flexible cognitive strategies during motor learning. PLoS Comput Biol. 2011;7: e1001096.

42. McDougle SD, Ivry RB, Taylor JA. Taking aim at the cognitive side of learning in sensorimotor adaptation tasks. Trends Cogn Sci. 2016;20: 535–544.

43. Bo J, Seidler RD. Visuospatial working memory capacity predicts the organization of acquired explicit motor sequences. J Neurophysiol. 2009;101: 3116–3125.

44. Anguera JA, Reuter-Lorenz PA, Willingham DT, Seidler RD. Contributions of spatial working memory to visuomotor learning. J Cogn Neurosci. 2010;22: 1917–1930.

45. Taylor JA, Thoroughman KA. Motor adaptation scaled by the difficulty of a secondary cognitive task. PLoS ONE. 2008;3: e2485.

46. Eversheim U. Evidence for processing stages in skill acquisition: A dual-task study. Learn Mem. 2001;8: 183–189.

47. Shadmehr R, Holcomb HH. Neural correlates of motor memory consolidation. Science. 1997;277: 821–825.

48. Anguera JA, Russell CA, Noll DC, Seidler RD. Neural correlates associated with intermanual transfer of sensorimotor adaptation. Brain Res. 2007;1185: 136–151.

49. Seidler RD, Bo J, Anguera JA. Neurocognitive contributions to motor skill learning: The role of working memory. J Mot Behav. 2012;44: 445–453.

50. Sweller J. Cognitive load during problem solving: Effects on learning. Cogn Sci. 1988;12: 257–285.

51. Sweller J, Merrienboer JJG van, Paas FGWC. Cognitive architecture and instructional design. Educ Psychol Rev. 1998;10: 251–296.

52. Noto CT, Watanabe S, Fuchs AF. Characteristics of simian adaptation fields produced by behavioral changes in saccade size and direction. J Neurophysiol. 1999;81: 2798–2813.

53. Ethier V, Zee DS, Shadmehr R. Changes in control of saccades during gain adaptation. J Neurosci. 2008;28: 13929–13937.

54. Panouillères M, Weiss T, Urquizar C, Salemme R, Munoz DP, Pélisson D. Behavioral evidence of separate adaptation mechanisms controlling saccade amplitude lengthening and shortening. J Neurophysiol. 2009;101: 1550–1559.

55. Zimmermann E, Lappe M. Motor signals in visual localization. J Vis. 2010;

56. Schnier F, Lappe M. Differences in intersaccadic adaptation transfer between inward and outward adaptation. J Neurophysiol. 2011;106: 1399–1410.

57. Catz N, Dicke PW, Thier P. Cerebellar-dependent motor learning is based on pruning a Purkinje cell population response. Proc Natl Acad Sci USA. 2008;105: 7309–7314.

58. Panouillères M, Neggers SFW, Gutteling TP, Salemme R, Stigchel S van der, Geest JN van der, et al. Transcranial magnetic stimulation and motor plasticity in human lateral cerebellum: Dual effect on saccadic adaptation. Hum Brain Mapp. 2012;33: 1512–1525.

59. Panouillères MTN, Miall RC, Jenkinson N. The role of the posterior cerebellum in saccadic adaptation: A transcranial direct current stimulation study. J Neurosci. 2015;35: 5471–5479.

60. Golla H, Tziridis K, Haarmeier T, Catz N, Barash S, Thier P. Reduced saccadic resilience and impaired saccadic adaptation due to cerebellar disease. Eur J Neurosci. 2008;27: 132–144.

61. Barash S, Melikyan A, Sivakov A, Zhang M, Glickstein M, Thier P. Saccadic dysmetria and adaptation after lesions of the cerebellar cortex. J Neurosci. 1999;19: 10931–10939.

62. Rolfs M, Knapen T, Cavanagh P. Global saccadic adaptation. Vision Res. 2010;50: 1882–1890.

63. Engbert R, Mergenthaler K. Microsaccades are triggered by low retinal image slip. Proc Natl Acad Sci USA. 2006;103: 7192–7197.

64. Hopp JJ, Fuchs AF. The characteristics and neuronal substrate of saccadic eye movement plasticity. Prog Neurobiol. 2004;72: 27–53.

65. Taylor JA, Ivry RB. Cerebellar and prefrontal cortex contributions to adaptation, strategies, and reinforcement learning. Prog Brain Res. 2014;210: 217–253.

66. Bender J, Tark K-J, Reuter B, Kathmann N, Curtis CE. Differential roles of the frontal and parietal cortices in the control of saccades. Brain Cognition. 2013;83: 1–9.

67. Duncan J. The multiple-demand (MD) system of the primate brain: Mental programs for intelligent behaviour. Trends Cogn Sci. 2010;14: 172–179.

68. Hutton SB, Ettinger U. The antisaccade task as a research tool in psychopathology: a critical review. Psychophysiology. 2006;43: 302–313.

69. Benson BL, Anguera JA, Seidler RD. A spatial explicit strategy reduces error but interferes with sensorimotor adaptation. J Neurophysiol. 2011;105: 2843–2851.

70. Craik FI, Salthouse TA. Handbook of Aging and Cognition II. Mahwah: Lawrence Erlbaum; 2000.

71. Raz N, Gunning FM, Head D, Dupuis JH, McQuain J, Briggs SD, et al. Selective aging of the human cerebral cortex observed in vivo: Differential vulnerability of the prefrontal gray matter. Cereb Cortex. 1997;7: 268–282.

72. Raz N, Lindenberger U, Rodrigue KM, Kennedy KM, Head D, Williamson A, et al. Regional brain changes in aging healthy adults: General trends, individual differences and modifiers. Cereb Cortex. 2005;15: 1676–1689.

73. Allen JS, Bruss J, Brown CK, Damasio H. Normal neuroanatomical variation due to age: The major lobes and a parcellation of the temporal region. Neurobiol Aging. 2005;26: 1245–1260.

74. Bock O. Components of sensorimotor adaptation in young and elderly subjects. Exp Brain Res. 2005;160: 259–263.

75. Bock O, Girgenrath M. Relationship between sensorimotor adaptation and cognitive functions in younger and older subjects. Exp Brain Res. 2006;169: 400–406.

76. Fernández-Ruiz J, Hall C, Vergara P, Díiaz R. Prism adaptation in normal aging: slower adaptation rate and larger aftereffect. Brain Res Cogn Brain Res. 2000;9: 223–226.

77. Buch ER, Young S, Contreras-Vidal JL. Visuomotor adaptation in normal aging. Learn Mem. 2003;10: 55–63.

78. McNay EC, Willingham DB. Deficit in learning of a motor skill requiring strategy, but not of perceptuomotor recalibration, with aging. Learn Mem. 1998;4: 411–420.

79. Heuer H, Hegele M, Sülzenbrück S. Implicit and explicit adjustments to extrinsic visuo-motor transformations and their age-related changes. Hum Mov Sci. 2011;30: 916–930.

80. Heuer H, Hegele M. Generalization of implicit and explicit adjustments to visuomotor rotations across the workspace in younger and older adults. J Neurophysiol. 2011;106: 2078–2085.

81. Anguera JA, Reuter-Lorenz PA, Willingham DT, Seidler RD. Failure to engage spatial working memory contributes to age-related declines in visuomotor learning. J Cognitive Neurosci. 2011;23: 11–25.

82. Langan J, Seidler RD. Age differences in spatial working memory contributions to visuomotor adaptation and transfer. Behav Brain Res. 2011;225: 160–168.

83. Munoz DP, Broughton JR, Goldring JE, Armstrong IT. Age-related performance of human subjects on saccadic eye movement tasks. Exp Brain Res. 1998;121: 391–400.

84. Fischer B, Biscaldi M, Gezeck S. On the development of voluntary and reflexive components in human saccade generation. Brain Res. 1997;754: 285–297.

85. Fukushima J, Hatta T, Fukushima K. Development of voluntary control of saccadic eye movements: I. Age-related changes in normal children. Brain and Development. 2000;22: 173–180.

